# Parallel evolution in mosquito vectors – a duplicated esterase locus is associated with resistance to pirimiphos-methyl in *An. gambiae*

**DOI:** 10.1101/2024.02.01.578361

**Authors:** Sanjay C. Nagi, Eric R. Lucas, Alexander Egyir-Yawson, John Essandoh, Samuel Dadzie, Joseph Chabi, Luc S. Djogbénou, Adandé A. Medjigbodo, Constant V. Edi, Guillaume K. Ketoh, Benjamin G. Koudou, Faisal Ashraf, Chris S. Clarkson, Alistair Miles, David Weetman, Martin J. Donnelly

**Author notes:** Corresponding author: Liverpool Tropical School of Medicine, Pembroke Place, Liverpool, L3 5QA, UK.

## Abstract

The primary control methods for the African malaria mosquito, *Anopheles gambiae*, are based on insecticidal interventions. Emerging resistance to these compounds is therefore of major concern to malaria control programmes. The organophosphate, pirimiphos-methyl, is a relatively new chemical in the vector control armoury but is now widely used in indoor residual spray campaigns. Whilst generally effective, phenotypic resistance has developed in some areas in malaria vectors. Here, we used a population genomic approach to identify novel mechanisms of resistance to pirimiphos-methyl in *Anopheles gambiae s*.*l* mosquitoes. In multiple populations, we found large and repeated signals of selection at a locus containing a cluster of detoxification enzymes, some of whose orthologs are known to confer resistance to organophosphates in *Culex pipiens*. Close examination revealed a pair of alpha-esterases, *Coeae1f* and *Coeae2f*, and a complex and diverse pattern of haplotypes under selection in *An. gambiae, An. coluzzii* and *An. arabiensis*. As in *Cx. pipiens*, copy number variation seems to play a role in the evolution of insecticide resistance at this locus. We used diplotype clustering to examine whether these signals arise from parallel evolution or adaptive introgression. Using whole-genome sequenced phenotyped samples, we found that in West Africa, a copy number variant in *Anopheles gambiae* is associated with resistance to pirimiphos-methyl. Overall, we demonstrate a striking example of contemporary parallel evolution which has important implications for malaria control programmes.

## Introduction

The spread of organophosphate resistance in the common house mosquito, *Culex pipiens*, is a textbook example of contemporary evolution in response to anthropogenic pressures (Raymond *et al*., 1998). In this species, mutations around two alpha-esterases enhanced the ability of the mosquito to detoxify organophosphate insecticides used in larviciding campaigns (Guillemaud *et al*., 1997). This locus was termed *Ester*, with independent gene duplications and transposable element insertions at the *Est2* and *Est3* genes resulting in at least 16 distinct haplotypes across the mosquitoes’ worldwide range (Raymond *et al*., 1996).

In the major malaria vector, *Anopheles gambiae*, resistance to organophosphates has historically been associated primarily with the *Ace1* locus, the target of organophosphate and carbamate insecticides. At this locus, a complex combination of heterogeneous or homogeneous gene duplications and the *Ace1*-G119S non-synonymous mutation, confer varying levels of resistance and fitness costs (Edi *et al*., 2014; Assogba *et al*., 2018; Grau-Bové *et al*., 2020). In recent years, organophosphates have been used increasingly for malaria vector control, primarily in indoor-residual spraying (IRS) campaigns as the formulation Actellic-300CS, in which pirimiphos-methyl is the active ingredient (Oxborough, 2016). Resistance to the compound has been slow to arise, and the *Ace1* locus remains the only validated resistance marker (Grau-Bové *et al*., 2020). Importantly, resistance to compounds in indoor residual spray formulations has been previously associated with IRS failure (Epstein *et al*., 2022).

Given the recent widespread increase in the use of organophosphates in malaria vector control, we investigate whether there is evidence for the evolution of novel organophosphate resistance mechanisms in *Anopheles gambiae*. Using data from the *Anopheles gambiae* 1000 genomes project (Miles *et al*., 2017; Clarkson *et al*., 2020), we found evidence of large, repeated signals of selection at the locus orthologous to the *Culex pipiens Ester* locus. We integrate expression data from studies across sub-Saharan Africa and perform an extensive analysis of this genomic region, which reveals that the *Coeae1f* and *Coeae2f* genes are an important resistance-associated locus. We find distinct copy number variants (CNVs) have arisen at this locus in both *An. gambiae* and *An. arabiensis* and that putative adaptive haplotypes have introgressed between *An. gambiae* and *An. coluzzii*. Lastly, we demonstrate that a CNV in *An. gambiae* is associated with resistance to the organophosphate pirimiphos-methyl. These data illustrate the importance of parallel evolution and introgression in the evolution of adaptively important traits in insect disease vectors.

## Results

### Data overview

We examined whole-genome sequence data from 2431 individual mosquitoes collected across eight countries of sub-Saharan Africa between the years 2012 to 2017 (Figure 1). These data are public through the MalariaGEN Vector Observatory (Clarkson *et al*., 2020) and GAARD projects (Lucas *et al*., 2023) (see methods for further details on sample collections and sequencing). A subset of 828 of these mosquitoes were phenotyped against the insecticides pirimiphos-methyl (n=347) or deltamethrin (n=481). To ensure strong differentiation between phenotypic groups, mosquitoes were classified into whether they were alive after exposure to a high dose of insecticide (resistant) or dead after exposure to a low dose (susceptible). Full details of the bioassay protocol are given in Lucas *et al*. 2023. For genome-wide selection scans and allele frequency estimates, we split the *An. gambiae* and *An. coluzzii* cohorts into early (2012-2015) and late (2017-2018) cohorts, to provide greater resolution to detect any low-frequency resistance mutants.

**Figure 1.**
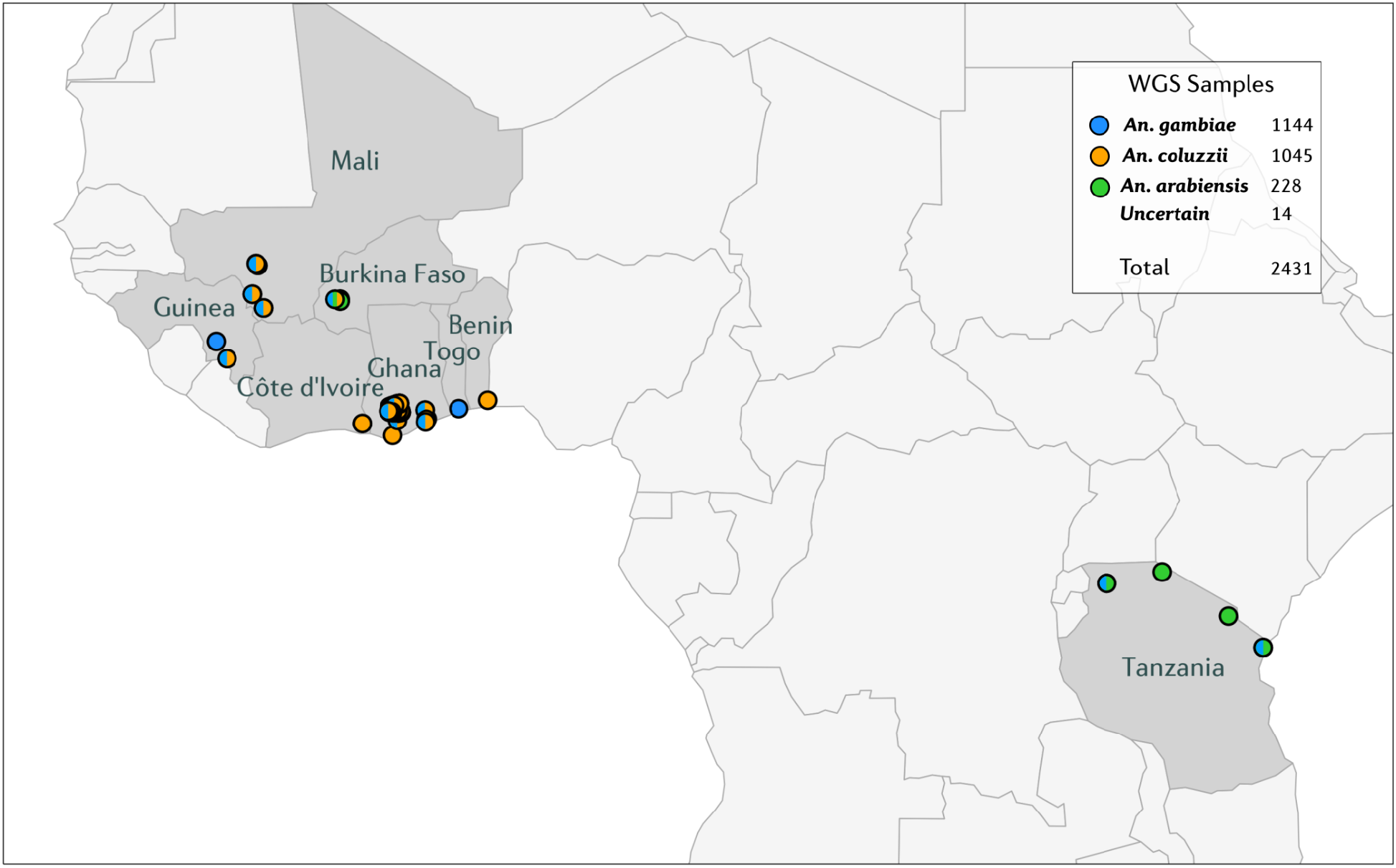
Map of sample sites for whole-genome sequenced individuals included in this study. Sample collection countries are highlighted in grey and collection sites as circles. Collection sites are coloured by species.

### A novel insecticide resistance locus

We investigated genome-wide signals of recent selection using the H12 statistic (Garud *et al*., 2015) in early (2012-2015) and late (2017-2018) cohorts of *An. gambiae, An. coluzzii*, and *An. arabiensis*, across all chromosomal arms. In addition to known selection signals at the *Vgsc* and *Rdl* resistance-associated loci, a novel, large signal of selection was observed on the 2L chromosomal arm in multiple populations, at (≅28.545 Mb, Figure 2A). This signal(s) was common in *An. gambiae* from West Africa and *An. arabiensis* from East Africa, but absent from *An. coluzzii*. The signals of selection are broad, with haplotype homozygosity inferred from the H12 signal extending beyond one megabase, suggesting that selection at the locus may have occurred relatively recently.

**Figure 2.**
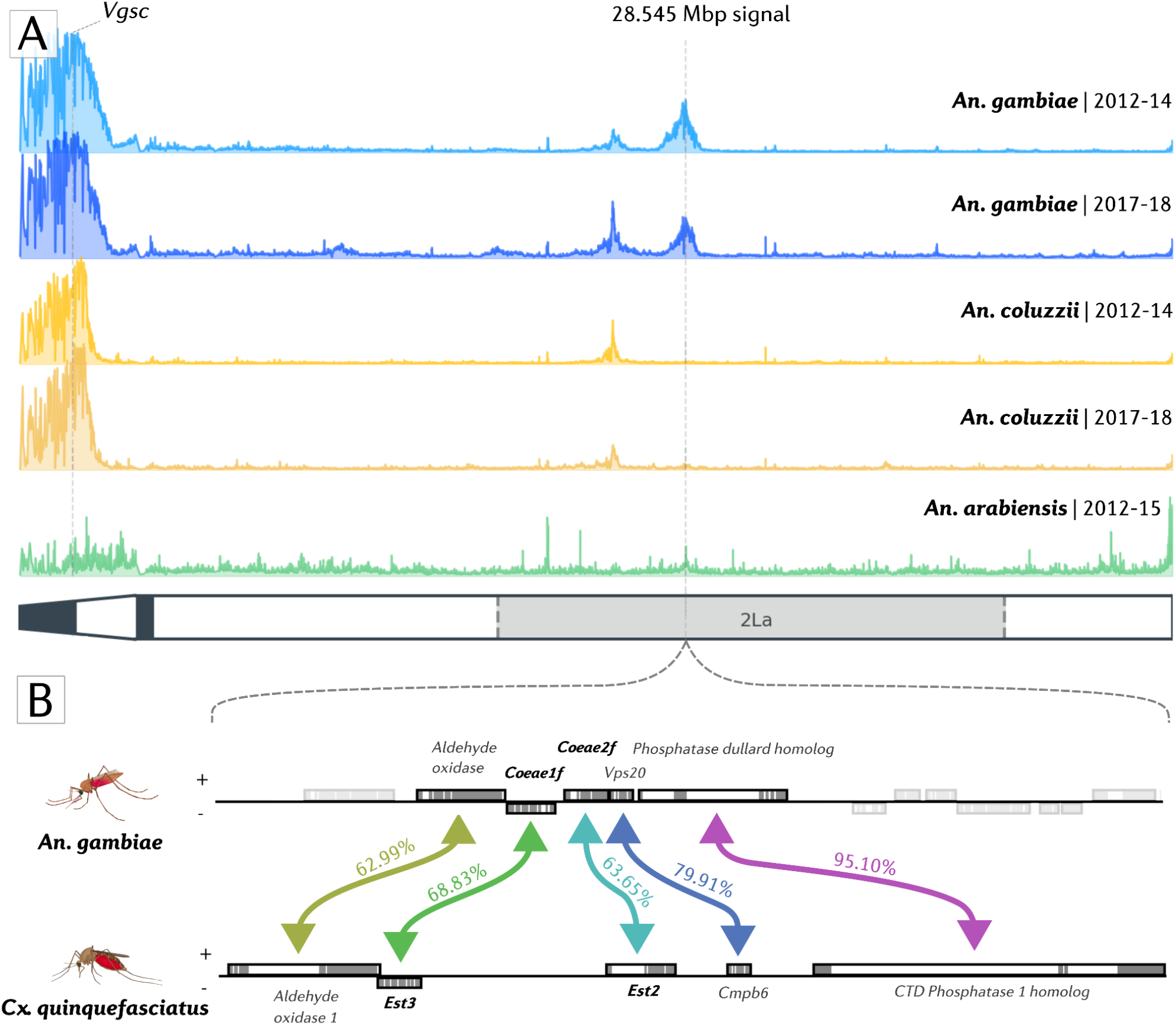
Selection scans and orthology of the alpha esterase cluster. A) H12 Genome-wide selection scans on the 2L chromosomal arm, in early and later cohorts of *An. gambiae* (blue), *An. coluzzii* (orange) and *An. arabiensis* (green). H12 was applied in 1500bp windows to phased, biallelic SNP haplotypes. A diagram of the 2L arm is shown, with heterochromatin coloured black and the 2La inversion shaded. The location of the *Coeae1f/2f* locus (28.545 Mbp) is highlighted with a dashed grey line. B) Orthology of *An. gambiae* (AgamP4, 2L:28,530,000-28,580,000) *Coeae1f/2f* locus with the *Cx. quinquefasciatus Est2* and *Est3* locus (JHB2020, CM027412.1:137,360,000-137,410,000). One-to-one orthologous pairs are highlighted with coloured arrows. Genes with orthologs not present within the 50Kb window are greyed out. BLASTp protein sequence similarity is labelled for each orthologous pair.

A closer examination of the region reveals several genes including two alpha-esterases, *Coeae1f* (AGAP006227) and *Coeae2f* (AGAP006228), that lie directly under the peak of the majority of the selection signals (Figure 2). These alpha-esterases sit on opposing strands in reverse orientation, 495 bases apart, and contain a differing number of exons (*Coeae1f:* 7; *Coeae2f: 4*). At the amino acid level, sequence similarity is 50.93% between the primary transcripts of the two genes. To identify relationships with known insecticide resistance genes in other species, we performed reciprocal blast searches with two other major vectors, *Aedes aegypti* and *Culex quinquefasciatus (*a member of the *Cx. pipiens* complex) (Supplementary Table 2). This revealed that the two carboxylesterases are one-to-one orthologs with the *Est3 (Coeae1f)* and *Est2 (Coeae2f)* carboxylesterases of *Cx. pipiens*, providing tentative evidence that *Coeae1f/2f* may be driving the signals of selection in *An. gambiae s*.*l*., given the confirmed role of *Est2 and Est3* in organophosphate resistance (Mouchès *et al*., 1986). Figure 2B displays a 50kb window in both the *An. gambiae* AgamP4 and *Cx. quinquefasciatus* JHB2020 genome assemblies, highlighting the orthology and synteny between the two species in this genomic region. *Coeae1f* shares 68.83% amino acid similarity with *Est3*, and *Coeae2f* shares 63.65% amino acid similarity with *Est2*. Despite other detoxification genes in the genomic region, we focus our analyses on the two alpha-esterases due to the strength of evidence in their favour.

Insecticide resistance is often associated with increased expression of insecticide-detoxifying genes, known as metabolic resistance (Ingham *et al*., 2018). Given that the two carboxylesterases are detoxification enzymes and that the locus has been associated with increased gene expression in *Cx. pipiens*, we postulated that they may also exhibit elevated expression in resistant strains of *An. gambiae s*.*l*. To determine whether there was any evidence for resistance-associated differential expression in *Coeae1f* and *Coeae2f*, we used the tool *AnoExpress*, which integrates gene expression data from 54 microarray (n=31) and RNA-Sequencing (n=23) experiments (Ingham *et al*., 2018; Nagi, 2023). Each of these experiments compared an insecticide-resistant strain of *An. gambiae s*.*l* to an insecticide-susceptible strain. Figure 3 displays log2 fold changes from each transcriptomic experiment, coloured by species.

**Figure 3.**
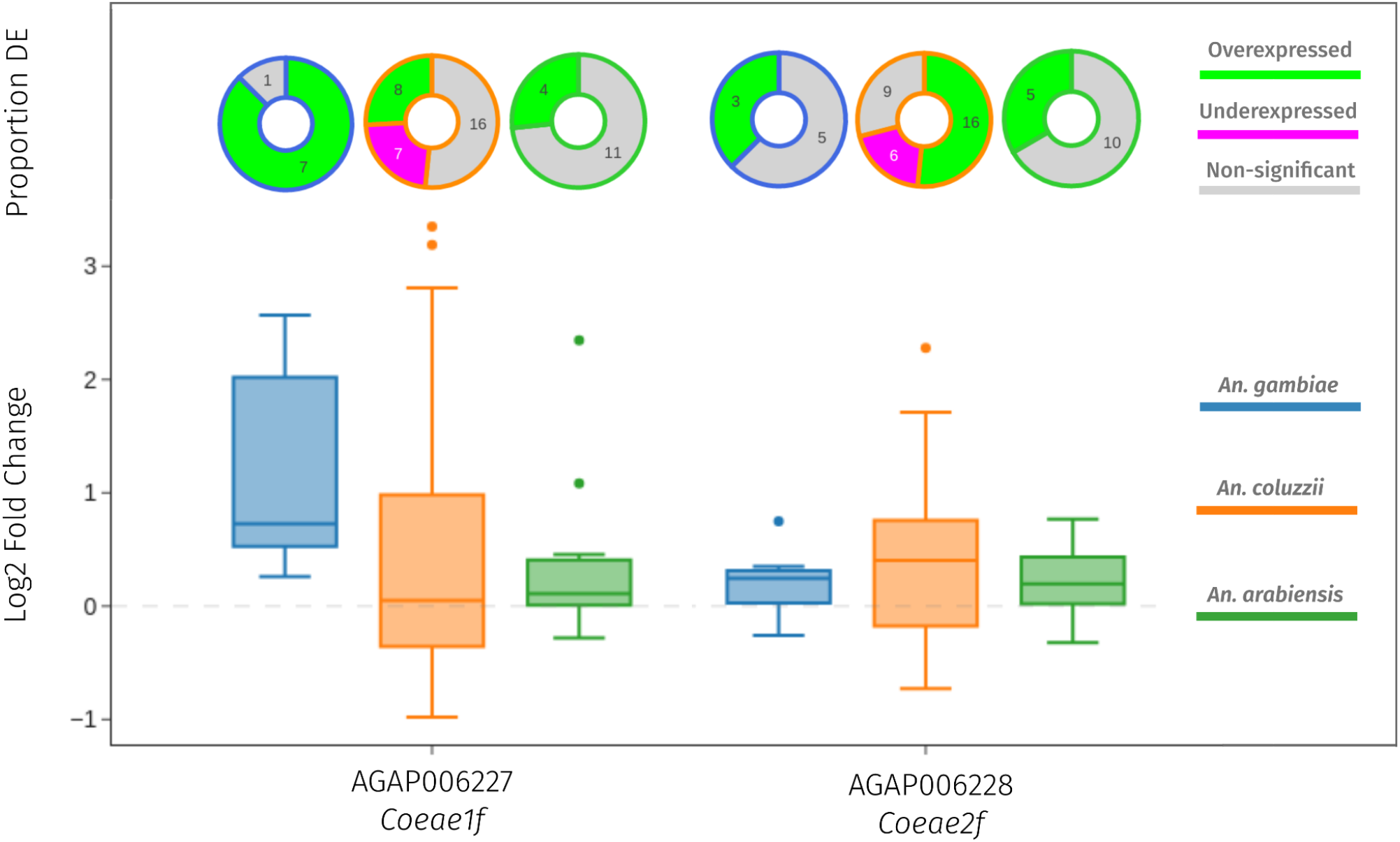
Gene expression data from AnoExpress, a meta-analysis of 54 transcriptomic studies into insecticide resistance in *An. gambiae* (blue), *An. coluzzii* (orange), and *An. arabiensis* (green). Bottom) A box plot of log2 fold changes in 54 transcriptomic experiments. Top) Donut charts showing the proportion of experiments which showed over-expression (green), under-expression (purple) or no differential expression (grey).

*Coeae1f* shows positive mean fold changes in *An. gambiae* (2.25), *An. coluzzii* (1.45) *and An. arabiensis* (1.26), across all microarray and RNA-Sequencing studies. It is very commonly over-expressed in *An. gambiae* (7/8 experiments), but less so in *An. coluzzii* (8/31) and *An. arabiensis* (4/15). *Coeae2f* also shows higher expression in resistant, compared to susceptible strains, with mean fold-changes of 1.15 in *An. gambiae*, 1.32 in *An. coluzzii*, and 1.16 in *An. arabiensis*.

Normalised read counts are high for both genes in the meta-analysis dataset (Supplementary Table 3), suggesting that they are highly expressed at a base level, and so could still contribute to the insecticide-resistant phenotype without large fold changes. Overall, the expression data is suggestive of a potential role in insecticide resistance, however, there is little spatial or temporal overlap between the regions in which we observe large selection signals and the transcriptomic data itself. In the only recent West African *An. gambiae* experiment (from Côte d’Ivoire), we observe a large fold-change of 5.93 for *Coeae1f*, from a region with established pirimiphos-methyl resistance, though it cannot be confirmed that this population contains haplotypes under selection in our data.

### Diplotype clustering of the *Coeae1f/2f* locus

To further explore patterns of selection at the *Coeae1f/2f* locus, we performed hierarchical clustering on diplotypes, to identify clusters of individuals under selection. A diplotype is a stretch of diploid genotypes, sometimes referred to as a multi-locus genotype. We used unphased diplotypes rather than haplotypes, to allow us to resolve multiallelic amino acid mutations and copy number variants, which are both typically challenging to phase onto haplotype scaffolds.

Figure 4 shows the results of this clustering, aligned with sample taxon, heterozygosity, the diplotype cluster a sample is assigned to, and the number of extra copies of *Coeae1f*. Supplementary Figure 3 contains the same plot for *Coeae2f*. Selective sweeps can be identified where there are multiple diplotypes (the leaves of the dendrogram) with zero or very low genetic distances between each other, and are highlighted by numbered diplotype cluster. To obtain clusters of closely related diplotype clusters, we cut the tree at a genetic distance of 0.04 and set a minimum cluster size of 40. Supplementary Table 4 contains a summary table of diplotype clusters and associated metadata. In diplotype cluster 1, which is primarily *An. coluzzii* individuals, we also find two *An. gambiae*, suggestive of potential adaptive introgression between species. There is no other evidence of adaptive introgression in other diplotype clusters.

**Figure 4.**
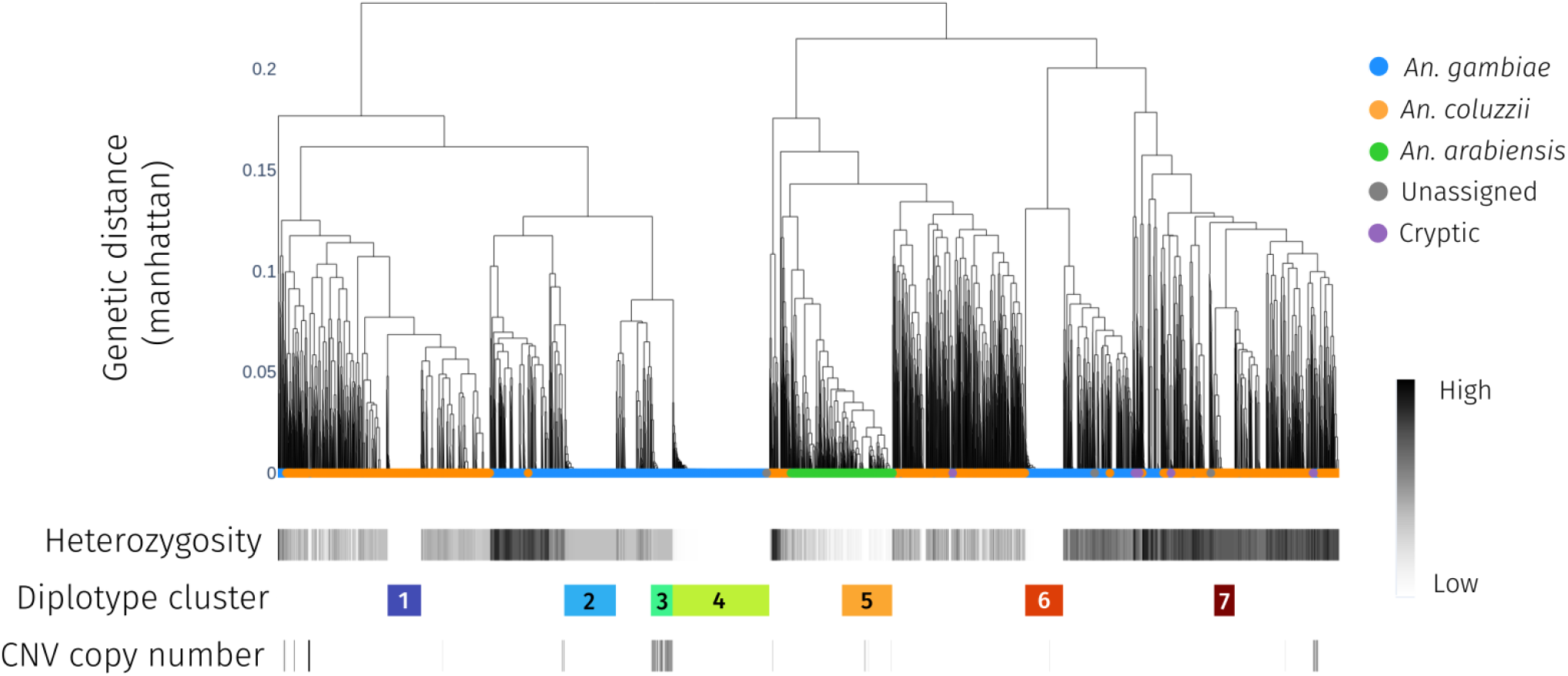
Diplotype clustering over the *Coeaexf* region. We calculate pairwise distance between diplotypes spanning the start of Coeae1f to the end of Coeae2f. Each column is a Diplotype ordered by the dendrogram with hierarchical clustering, using genetic distance based on cityblock (Manhattan) distance and complete linkage. Clusters with low sample heterozygosity or inter-sample distances of zero are indicative of a selective sweep. CNVs; A CNV is strongly associated with a sweep found in West African *An. gambiae (*cluster 3), and another one weakly associated with a sweep in Tanzanian *An. arabiensis* (cluster 5)..

We investigated amino acid variation at the *Coeae1f* and *Coeae2f* genes by plotting amino acid variation alongside the diplotype clustering dendrograms (Supplementary Figures 1 and 2), to identify potential mutants driving selective sweeps. The majority of amino acid mutations found on selective sweep diplotypes are also found at moderate to high frequencies in wild-type individuals and are therefore not obvious candidates to be causal mutations. There are, however, some exceptions - *Coeae1f-*E477V is relatively rare outside of swept diplotypes, as are *Coeae1f-*Q129E, *Coeae1f-*D463N, *Coeae2f-S*357N and *Coeae2f-*V516L.

### Copy number variation

In *Cx. pipiens*, copy number variants at the *ester* locus that cover one or both of the *Coeae1f* and *Coeae2f* orthologs have spread around the world (Raymond *et al*., 1991). Given the gene expression data and the presence of copy number variation at this locus in *Cx. pipiens*, we speculated whether copy number variants might exist at these loci in *An. gambiae s*.*l*. There is increasing evidence of CNVs and metabolic resistance associations (Lucas *et al*., 2019; Njoroge *et al*., 2022; Lucas *et al*., 2023). To identify CNVs, we calculated the copy number at each gene in the region and then computed the frequency of amplifications or deletions in the dataset.

The combined copy number and diplotype clustering analysis suggested that CNVs were associated with two selective sweeps, one strongly associated with diplotype cluster 3, found in *An. gambiae*, and one very weakly with diplotype cluster 5, found in *An. arabiensis* (Figure 4). In the *An. gambiae* CNV-associated diplotype cluster, heterozygosity is elevated, suggesting that cluster contains individuals heterozygous for the CNV and a distinct selective sweep haplotype. We found the *An. gambiae* CNV only in West African populations of *An. gambiae*, and we called this duplication *Coeaexf*-Dup1. The CNV in *An. arabiensis* was found in a cohort from Tanzania, we call this *Coeaexf*-Dup2. Supplementary Figure 3B shows the copy number of *Coeae1f* for each diplotype cluster.

We then examined the genomic span of each copy number variant. Supplementary Figure 4 shows sequence coverage data in 300bp windows for two representative individuals from Ghana and Tanzania. Both CNVs cover the two carboxylesterases *Coeae1f* and *Coeae2f*, and the *Coeaexf*-Dup1 CNV in Ghana and Togo only covers these two genes - also amplifying a truncated version of a putative aldehyde oxidase, AGAP006226 (Figure 5A). *Coeaexf*-Dup2 is much larger, and spans ten genes in total. Figure 5B shows a summary of the frequencies of CNVs at the locus in each cohort. No other CNVs were noted covering detoxification genes at the locus.

**Figure 5.**
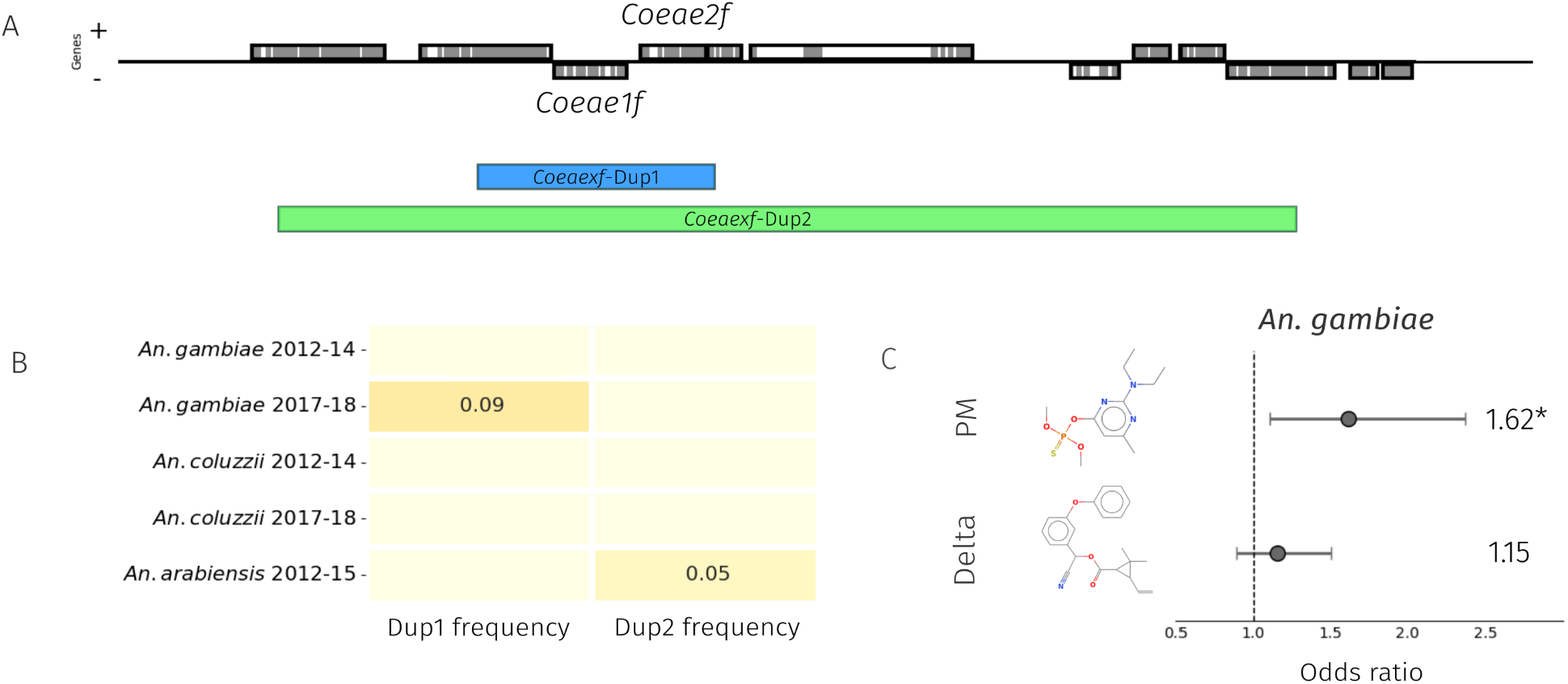
A schematic of the *Coeae1f/2f* locus, with associated frequency and genotype:phenotype associations. Frequencies of the presence of at least one Dup allele in our cohorts (B). A forest plot showing odds ratios from the binomial GLM assessing the relationship between *Coeae1f* copy number and survival to either pirimiphos-methyl or deltamethrin.

In our data, we find that individuals positive for *Coeaexf-*Dup1 have a maximum copy number of six, whereas those positive for *Coeaexf-*Dup2 have a maximum copy number of four.

### A CNV at *Coeae1f/Coeae2f* is associated with resistance to pirimiphos-methyl in West African *An. gambiae*

We hypothesised that copy number variation at *Coeae1f/2f* may confer resistance to organophosphate insecticides used in vector control. We specifically tested for associations against the active ingredient of Actellic-300CS, pirimiphos-methyl, and the pyrethroid deltamethrin. Deltamethrin is the most widely used insecticide in long-lasting insecticide-treated nets (LLINs), and due to the ubiquitous use of pyrethroids, resistance has spread through sub-Saharan Africa over the past two-decades.

To determine whether CNVs in *Coeae1f* and *Coeae2f* are associated with resistance to insecticides, we took a subset of samples from our *An. gambiae* cohorts which were phenotyped by WHO tube assays to either pirimiphos-methyl or deltamethrin. Details of this sample set have been published previously (Lucas *et al*., 2023). Reliable CNV data were available from 463 *An. gambiae* samples from four locations. We found CNVs in *Coeae1f* in 28 out of 146 samples (19%) from Baguida (Togo) and 23 out of 160 (14%) samples from Obuasi (Ghana), but such CNVs were absent from all 37 samples in Aboisso (Côte d’Ivoire) and 120 samples in Madina (Ghana). We tested for a significant association with phenotype in both of these populations and included CNVs in *Ace1* in the model due to their known association with pirimiphos-methyl resistance. When considering each population separately, we found a significant association with PM resistance in Obuasi (P = 0.034, OR = 1.76, 1.04-3.4) and marginally non-significant in Baguida (P = 0.077). When combining the two locations and including location as a random effect in the model, significance was increased compared to either site alone (P = 0.007, OR = 1.62, 1.1-2.36). In contrast, as expected, we found no significant association of *Coeae1f* CNVs with resistance to deltamethrin.

### CNV diagnostics

To track these CNVs in time and space, we have developed PCR primers for use in standard genotyping PCRs, which involve three different primers for each duplication. A forward and reverse primer combine to produce a band when the duplication is present. There is then also a control primer, which combines with either the forward or reverse primer to produce a control band, which should be present in all samples regardless of genotype. Primer sequences can be found in Supplementary Text 1. We tested these primers on a set of 20 samples which were either positive or negative for *Coeaexf-*Dup1 from West Africa, or positive or negative for *Coeaexf-*Dup2 from Tanzania. Concordance was 100% (Supplementary Figure 5).

## Discussion

In the common house mosquito, *Culex pipiens*, the *ester* locus, orthologous to the *Coeae1f/2f* locus, is a textbook example of contemporary anthropogenic selection (Raymond *et al*., 1998). Through exposure to toxic organophosphate insecticides, at least 16 haplotypes spread throughout the range of the mosquito, competing with each other and providing varying combinations of insecticide resistance and fitness cost. Many of these haplotypes are associated with gene amplifications around the two alpha-esterases.

In this study, we used whole-genome sequence data from 2431 individual mosquitoes to demonstrate parallel evolution at the orthologous locus in malaria mosquitoes. There is evidence of intense selection acting upon the locus in *An. gambiae, An. coluzzii, and An. arabiensis*, with multiple haplotypes rising to high frequencies, some of which harbour CNVs. We demonstrate that in *An. gambiae* gene amplifications at this locus are associated with resistance to the organophosphate pirimiphos-methyl. Some similarities between species are striking - in *Cx. pipiens* there is a gene amplification allele which, as with the *Coeaexf*-Dup1 allele in *An. gambiae*, covers both alpha esterases and a partial copy of the neighbouring aldehyde oxidase gene (Buss *et al*., 2004). Five of the seven diplotype clusters reported in this study do not contain copy number variants, and it is not clear what causal variants are driving these selective sweeps; amino acid mutations are a possibility, however, genomic insertions, which are commonly involved in insecticide resistance, could also be important.

As well as evidence of parallel evolution, the study also provides important information for malaria control programmes. We reveal a novel locus in *An. gambiae* which contributes to resistance to the organophosphate pirimiphos-methyl. Pirimiphos methyl, formulated as Actellic CS300, is widely used in indoor residual spraying (IRS) campaigns throughout sub-Saharan Africa. Before this study, only one locus in *An. gambiae s*.*l* had been associated with resistance to this compound - *Ace1*, which is the target site of organophosphate and carbamate insecticides. Unlike LLINs which still provide a physical barrier and so protect even against insecticide-resistant vectors, IRS is arguably more prone to resistance-mediated failure. Examples of this come from Kwazulu-Natal, where pyrethroid-resistant *Anopheles funestus* caused a malaria outbreak after a change in IRS product from DDT to pyrethroids (Maharaj *et al*., 2005) and more recently, from IRS campaigns in Uganda (Epstein *et al*., 2022). It is therefore even more important to detect emerging mechanisms of resistance early, so that insecticide resistance management (IRM) practices can be employed, such as rotating out the insecticide in favour of the new IRS active ingredients such as clothianidin, broflanilide and chlorfenapyr. The discoveries of mutations at both the *Ace1* and *Coeaexf* loci in *An. gambiae* mosquitoes from West Africa are important, and these mutations should be monitored as part of insecticide resistance management campaigns. We provide a CNV diagnostic PCR assay allowing researchers to monitor these variants in time and space. We stress that the discovery of this novel variant does not necessarily mean pirimiphos-methyl is no longer effective, but that this locus should be incorporated in future surveillance activities.

## Methods

### Data collection

We used a subset of whole genome sequence variation data from phase 3 of the *Anopheles* 1000 genomes project and a recent GWAS study in West Africa (Clarkson *et al*., 2020; Lucas *et al*., 2023). The data contained 2431 individual *Anopheles* mosquitoes, of which there were 1144 *An. gambiae*, 1045 *An. coluzzii*, 228 *An. arabiensis*, 11 of a cryptic taxon termed gcx3, and three unassigned individuals. Specimens were all collected between 2013 and 2017. Sample provenance and methods for genome sequencing and analysis are described elsewhere (Clarkson *et al*., 2020; Lucas *et al*., 2023).

### Genome-wide selection scans

We first performed genome-wide selection scans with the H12 statistic (Garud *et al*., 2015). This statistic captures the haplotype frequency spectrum of the two highest frequency haplotype clusters and is particularly powerful in detecting soft selective sweeps, or where there are multiple distinct haplotype clusters at the same locus. We first extracted phased biallelic haplotypes from the 2431 individual mosquitoes and split the populations into cohorts of West African (WA) *An. coluzzii*, WA *An. gambiae*, East African (EA) *An. gambiae and EA An. arabiensis*. We further split the WA cohorts into early (2012-2014) and later collections (2017-2018). We then took a random sample of 100 haplotypes from each cohort and used malariagen_data to compute H12 in 1500 SNP stepping windows.

### Orthology searches

We extracted the predicted amino acid sequences of AGAP006227 and AGAP006228 from Vectorbase. We used diamond (Buchfink *et al*., 2015) to perform sequence alignment with the *An. gambiae* PEST, *Cx. quinquefasciatus* JHB2020, and *Ae*.*aegypti* LVP_AGWG reference genomes (Vectorbase, version 66).

### Gene expression data

We used the Python package *AnoExpress* (Nagi, 2023) to load and visualise gene expression data. *AnoExpress* contains gene expression data from microarray and RNA-sequencing studies into insecticide resistance in *An. gambiae s*.*l* and *An. funestus*. RNA-Sequencing read count data were retrieved directly from authors and analysed with DESeq2 v1.26 (Love *et al*., 2014).

### Diplotype clustering

We extracted diplotypes from the start of *Coeae1f* to the end of the *Coeae2f* gene (2L:28,545,396-28,550,748) and clustered the diplotypes with complete-linkage hierarchical clustering. We first transform diplotypes into an array of allele counts, that is, for each site, an array of size 4, one for each possible allele. For each site, we then find the pairwise difference between this allele counts array for each individual in the pair. We then use city-block (Manhattan) distance as the distance metric and complete linkage. We also determined which non-synonymous variants were present on each diplotype and plotted this data alongside sample heterozygosity over the region, and the number of extra copies of *Coeae1f* (AGAP006227) and *Coeae2f* (AGAP006228).

### CNV association tests

Using copy number estimated for 828 phenotyped individuals from West Africa (Lucas *et al*., 2023) we performed CNV association tests. These individuals were phenotyped against either Pirimiphos-methyl or Deltamethrin, using WHO tube bioassays (WHO, 2016). We excluded samples with high variance in coverage, leaving 463 samples with reliable CNV calls. We first estimated the copy number at *Coeae1f* and converted this to a binary variable, whether an individual has at least one amplification. We performed CNV association tests for each location with a binomial GLM with logit link function using CNV status and *Ace1* genotype as predictor variables and phenotype (dead or alive) as the response variable. GLMs were performed in R v4 (R Core Team, 2018) and the inclusion of location as a random effect was implemented using the package glmmTMB (Brooks *et al*., 2017).

### Code availability

Code used to analyse the data are available in the GitHub repository https://github.com/sanjaynagi/coeaexf. All sequencing, alignment, SNP and CNV calling were carried out as part of the Anopheles gambiae 1000 genomes project v3.2 (https://www.malariagen.net/data).

## Acknowledgements

This work was supported by the National Institute of Allergy and Infectious Diseases (NIAID R01-AI116811 to M.J.D. and D.W.) and the Medical Research Council (MR/T001070/1 to M.J.D., D.W. and E.R.L., MR/ P02520X/1 to M.J.D. and D.W.). The latter grant is a UK-funded award and is part of the EDCTP2 programme supported by the European Union. M.J.D. is supported by a Royal Society Wolfson Fellowship (RSWF\FT \180003). SCN was funded by an MRC CASE studentship (MR/R015678/1).

This study was supported by the MalariaGEN Vector Observatory which is an international collaboration working to build capacity for malaria vector genomic research and surveillance, and involves contributions by the following institutions and teams. Wellcome Sanger Institute: Lee Hart, Kelly Bennett, Anastasia Hernandez-Koutoucheva, Jon Brenas, Menelaos Ioannidis, Chris Clarkson, Alistair Miles, Julia Jeans, Paballo Chauke, Victoria Simpson, Eleanor Drury, Osama Mayet, Sónia Gonçalves, Katherine Figueroa, Tom Madison, Kevin Howe, Mara Lawniczak; Liverpool School of Tropical Medicine: Eric Lucas, Sanjay Nagi, Martin Donnelly; Broad Institute of Harvard and MIT: Jessica Way, George Grant; Pan-African Mosquito Control Association: Jane Mwangi, Edward Lukyamuzi, Sonia Barasa, Ibra Lujumba, Elijah Juma. The authors would like to thank the staff of the Wellcome Sanger Genomic Surveillance unit and the Wellcome Sanger Institute Sample Logistics, Sequencing and Informatics facilities for their contributions.

The MalariaGEN Vector Observatory is supported by funding awarded to Dominic Kwiatkowski and Mara Lawniczak from Wellcome (220540/Z/20/A, ‘Wellcome Sanger Institute Quinquennial Review 2021-2026’) and funding awarded to Dominic Kwiatkowski from the Bill and Melinda Gates Foundation (INV-001927). The Liverpool School of Tropical Medicine’s participation was supported by the National Institute of Allergy and Infectious Diseases ([NIAID] R01-AI116811), with additional support from the Medical Research Council (MR/P02520X/1). The latter grant is a UK-funded award and is part of the EDCTP2 programme supported by the European Union. Martin Donnelly is supported by a Royal Society Wolfson Fellowship (RSWF\FT\180003). The Pan-African Mosquito Control Association’s participation was funded by the Bill and Melinda Gates Foundation (INV-031595).

## Author contributions

SCN, MJD, ERL and DW designed the study; ERL described the CNV alleles, designed the CNV primers and performed the phenotype association analysis; SCN performed all other analyses; AEY, JE, SD, JC, LSD, AAM, CVE, GKK, BGK conducted field collections and phenotyping; FA tested diagnostic primers; SCN, MJD, DW and ERL wrote the manuscript with input from all authors. All authors approved the final version of the manuscript.

