## Supplementary Information for "Parallel evolution in mosquito vectors – a duplicated esterase locus is associated with resistance to pirimiphos-methyl in *An. gambiae*"

### ***Coeaexf*** **supplementary information**

**Supplementary Table 1.** Sample manifest

| country | year | arabiensis | coluzzii | gambiae | gcx3 | unassigned |
|---|---|---|---|---|---|---|
| **Benin** | 2017 | 0 | 90 | 0 | 0 | 0 |
| **Burkina Faso** | 2012 | 0 | 82 | 99 | 0 | 0 |
| **Burkina Faso** | 2014 | 3 | 53 | 46 | 0 | 0 |
| **Cote d'Ivoire** | 2017 | 0 | 1 | 36 | 0 | 1 |
| **Ghana** | 2012 | 0 | 64 | 36 | 0 | 0 |
| **Ghana** | 2017 | 0 | 0 | 398 | 0 | 0 |
| **Ghana** | 2018 | 0 | 690 | 63 | 0 | 0 |
| **Guinea** | 2012 | 0 | 11 | 125 | 0 | 0 |
| **Mali** | 2012 | 0 | 27 | 65 | 0 | 2 |
| **Mali** | 2014 | 0 | 27 | 33 | 0 | 0 |
| **Tanzania** | 2012 | 87 | 0 | 0 | 0 | 0 |
| **Tanzania** | 2013 | 1 | 0 | 32 | 10 | 0 |
| **Tanzania** | 2015 | 137 | 0 | 32 | 1 | 0 |
| **Togo** | 2017 | 0 | 0 | 179 | 0 | 0 |

#### **Supplementary Table 2.** Reciprocal best hits.

Aligning AGAP006227, AGAP006228 protein sequences against the *Culex quinquefasciatus* JHB2020 and Aedes aegypti reference genomes. For the *Culex* genes, we then align back to the *An. gambiae* PEST reference. Only the top five hits for each search are shown.

CQUJHB000812 = *Est3* , CQUJHB006176 = *Est2*

| query_seqid | reference_seqid | Protein identity % | length | mismatch | evalue |
|---|---|---|---|---|---|
| AGAP006228-RA | CQUJHB006176.P9455 | 64.2 | 542 | 190 | 1.76E-264 |
| AGAP006228-RA | CQUJHB000812.P1285 | 51 | 537 | 257 | 6.7E-199 |
| AGAP006228-RA | CQUJHB006077.P9314 | 36.1 | 529 | 309 | 3.05E-90 |
| AGAP006228-RA | CQUJHB012326.P19058 | 34.6 | 552 | 329 | 3.31E-85 |
| AGAP006228-RA | CQUJHB012326.P19057 | 34.6 | 552 | 329 | 4.27E-85 |
| AGAP006227-RA | CQUJHB000812.P1285 | 68.8 | 539 | 165 | 1.97E-292 |
| AGAP006227-RA | CQUJHB006176.P9455 | 50.7 | 534 | 258 | 4.74E-197 |
| AGAP006227-RA | CQUJHB006077.P9314 | 35 | 545 | 322 | 8.31E-91 |
| AGAP006227-RA | CQUJHB005542.P8540 | 34.1 | 557 | 321 | 3.61E-89 |
| AGAP006227-RA | CQUJHB012326.P19058 | 33.8 | 541 | 331 | 3.94E-88 |
| CQUJHB000812 | AGAP006227-PA | 68.8 | 539 | 165 | 3.59E-291 |
| CQUJHB000812 | AGAP006228-PA | 51 | 537 | 257 | 2.42E-202 |
| CQUJHB000812 | AGAP002391-PA | 35.5 | 552 | 324 | 8.14E-93 |
| CQUJHB000812 | AGAP006700-PA | 32.7 | 556 | 345 | 1.53E-84 |
| CQUJHB000812 | AGAP006727-PA | 34 | 550 | 330 | 1.11E-83 |
| CQUJHB006176 | AGAP006228-PA | 64.2 | 542 | 190 | 1.78E-267 |
| CQUJHB006176 | AGAP006227-PA | 50.7 | 534 | 258 | 2.42E-195 |
| CQUJHB006176 | AGAP002391-PA | 38.9 | 458 | 259 | 7.33E-84 |
| CQUJHB006176 | AGAP006700-PA | 33.1 | 553 | 343 | 7.07E-81 |
| CQUJHB006176 | AGAP006726-PA | 36.4 | 456 | 279 | 1.72E-79 |
| AGAP006228-RA | AAEL017071-PA | 64.4 | 534 | 189 | 3.29E-268 |
| AGAP006228-RA | AAEL010389-PA | 48.9 | 542 | 273 | 1.89E-192 |
| AGAP006228-RA | AAEL019679-PB | 33.8 | 553 | 332 | 2.38E-83 |
| AGAP006228-RA | AAEL019679-PC | 33.8 | 553 | 332 | 2.38E-83 |
| AGAP006228-RA | AAEL019679-PD | 33.8 | 553 | 332 | 2.38E-83 |
| AGAP006227-RA | AAEL010389-PA | 66.8 | 539 | 176 | 1.13E-283 |
| AGAP006227-RA | AAEL017071-PA | 52.2 | 533 | 249 | 3.94E-207 |
| AGAP006227-RA | AAEL019679-PB | 35.9 | 541 | 320 | 7.61E-95 |
| AGAP006227-RA | AAEL019679-PC | 35.9 | 541 | 320 | 7.61E-95 |
| AGAP006227-RA | AAEL019679-PD | 35.9 | 541 | 320 | 7.61E-95 |

**Supplementary Figure 1.** Amino acid variation at Coeae1f

NB. diplotype cluster assignments and colours may differ to the diplotype clustering plot in the main text, as the main text used the entire Coeaexf locus, rather than this plot which focuses on *Coeae1f*.

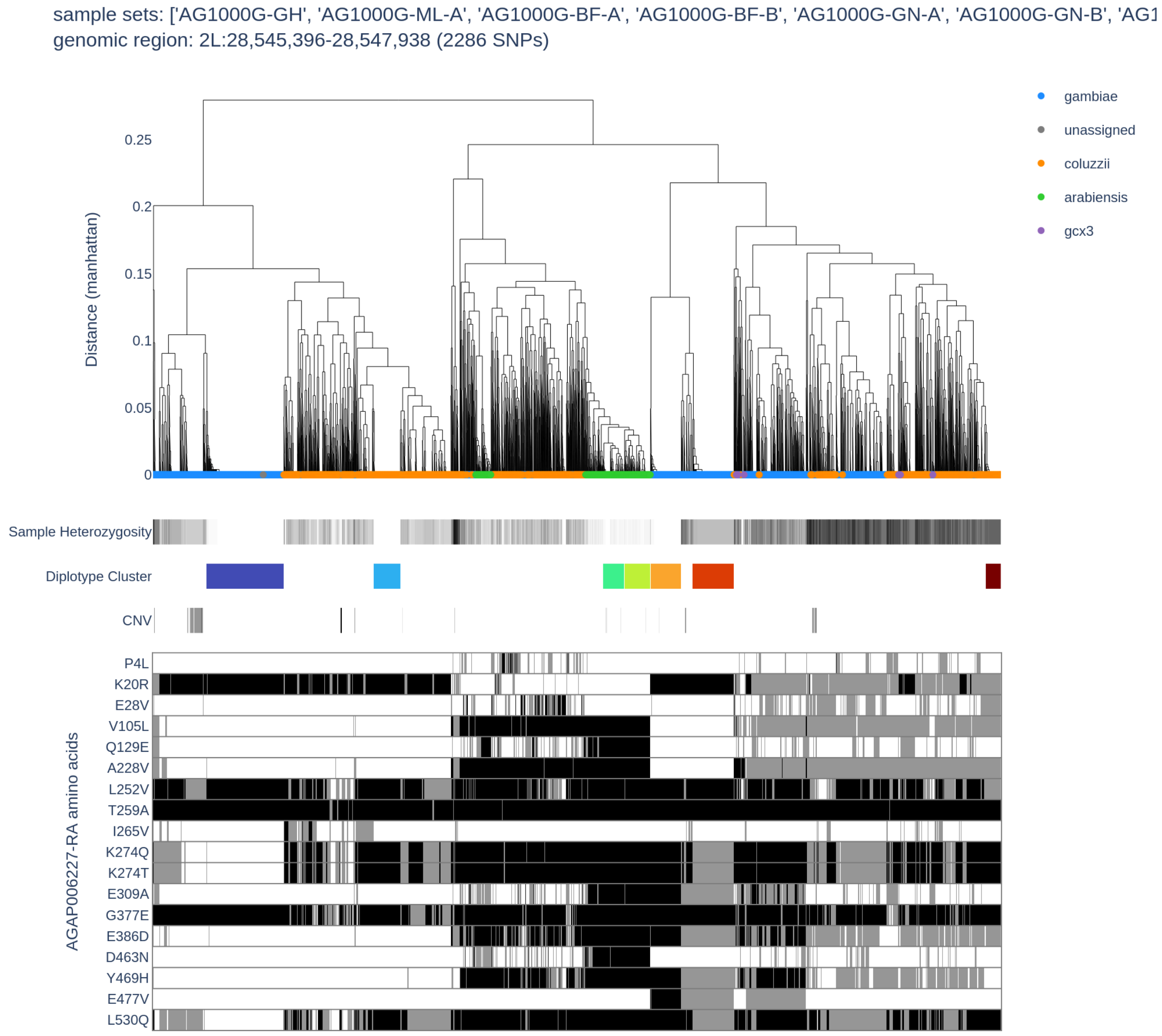


Supplementary Figure 1. **Diplotype clustering over the *Coeae1f* region.** We calculate pairwise distance between diplotypes spanning the start of Coeae1f to Coeae1f. Each column is a Diplotype ordered by the dendrogram with hierarchical clustering, using genetic distance based on cityblock (Manhattan) distance and complete linkage. Clusters with low sample heterozygosity or inter-sample distances of zero are indicative of a selective sweep. Amino acid variation is displayed alongside the dendrogram.

#### **Supplementary Figure 2.** Amino acid variation at Coeae2f

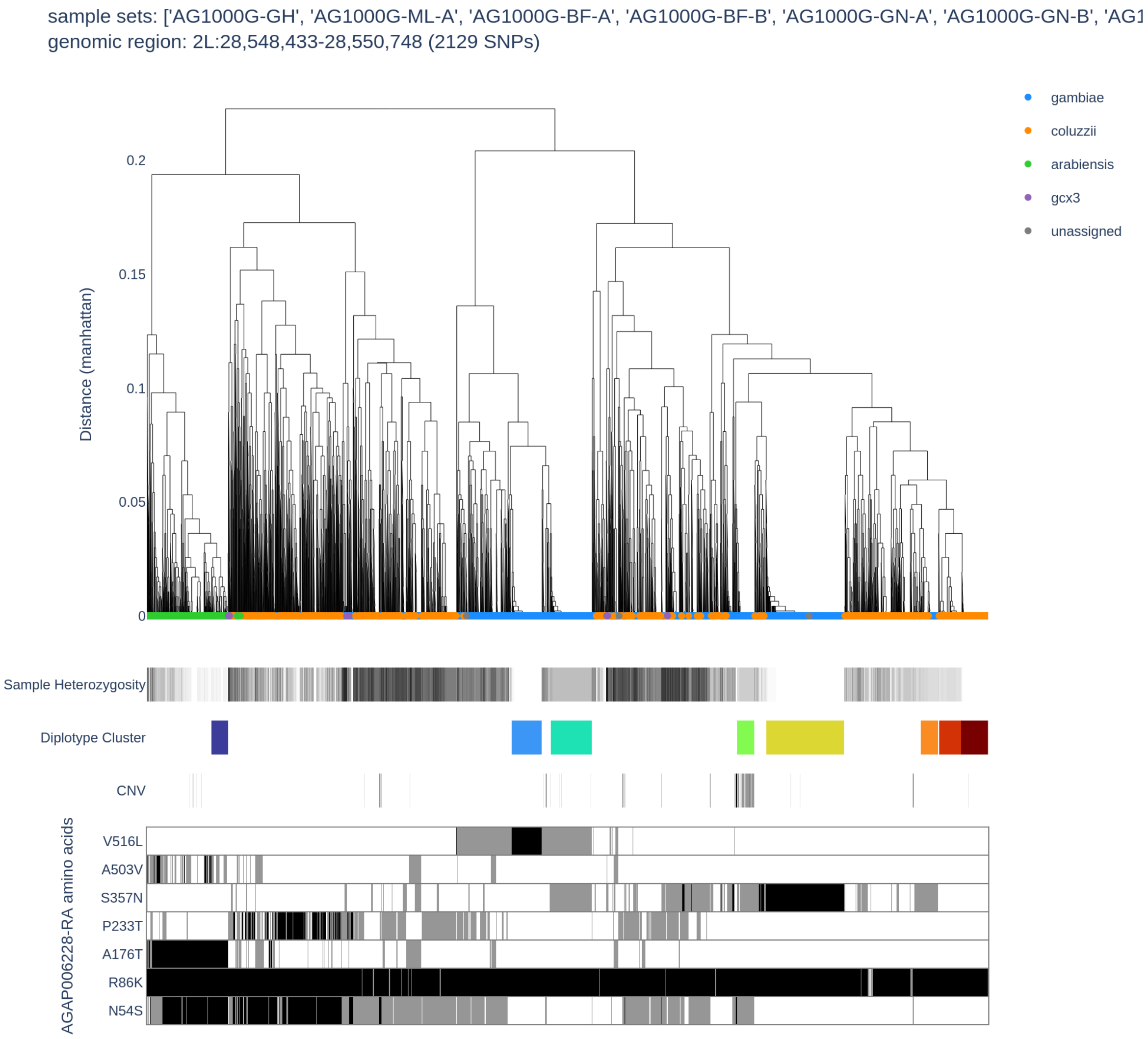


Supplementary Figure 1. **Diplotype clustering over the *Coeae2f* region.** We calculate pairwise distance between diplotypes spanning the start of Coeae2f to the end of Coeae2f. Each column is a Diplotype ordered by the dendrogram with hierarchical clustering, using genetic distance based on cityblock (Manhattan) distance and complete linkage. Clusters with low sample heterozygosity or inter-sample distances of zero are indicative of a selective sweep. Amino acid variation is displayed alongside the dendrogram.

**Supplementary Table 3.** Mean read counts of Coeae1/2f

| GeneID | mean counts |
|---|---|
| **AGAP006227** | 408.971150 |
| **AGAP006228** | 750.124806 |

#### **Supplementary Table 4.** Diplotype cluster summary

This table describes the number of individuals in our data who fall into each diplotype cluster, and the proportion of that cluster with CNV-positive individuals.

| cluster | arabiensis | coluzzii | gambiae | proportion_with_cnv |
|---|---|---|---|---|
| **WT** | 113 | 925 | 666 | 0.02 |
| **1** | 0 | 74 | 2 | 0 |
| **2** | 0 | 0 | 118 | 0.01 |
| **3** | 0 | 0 | 50 | 0.91 |
| **4** | 0 | 0 | 221 | 0 |
| **5** | 115 | 0 | 0 | 0.09 |
| **6** | 0 | 0 | 87 | 0.01 |
| **7** | 0 | 46 | 0 | 0 |

**Supplementary Figure 3.** Diplotype clusters and CNV Status/ Taxon

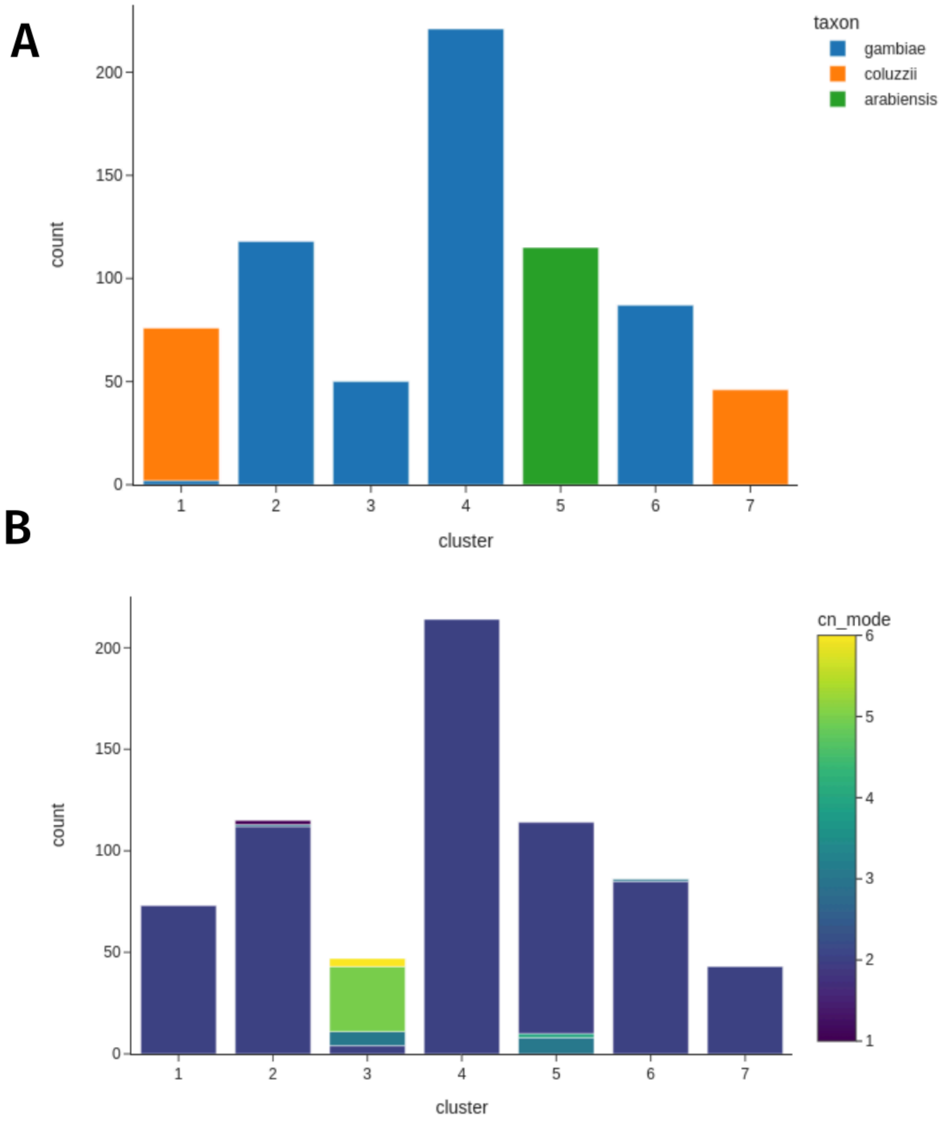


**Supplementary Figure 3A)** The number of individuals in each diplotype cluster, coloured by taxon assignment. 3B) The number of individuals in each diplotype cluster, coloured by copy number at Coeae1f.

**Supplementary Figure 4.** Example coverage traces for Dup1 and Dup2

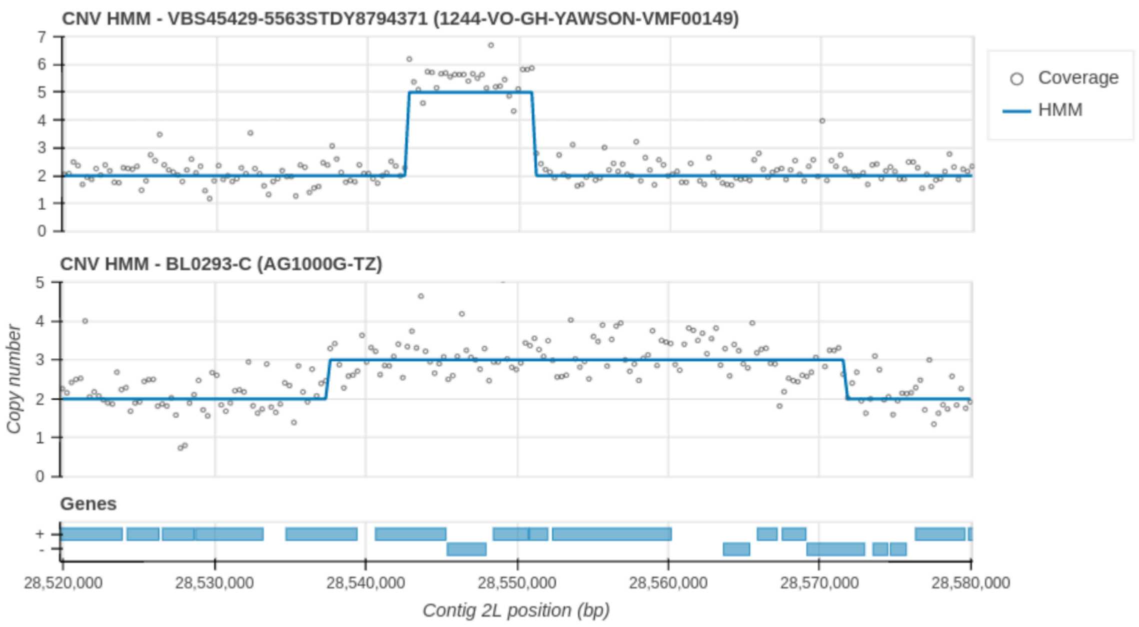


**Suppl. Figure 4.** Example coverage traces for Dup1 (upper, *An. gambiae female* collected in Obuasi, Ghana) and Dup2 (lower, *An. arabiensis female* from Moshi, Tanzania). Coverage is calculated in 300 Bp stepping windows, and the line represents the HMM prediction of copy number state described in (Lucas *et al.*, 2019).

#### **Supplementary Text 1** - CNV primer diagnostic protocols

| Primer name | Sequence |
|---|---|
| Coeaexf_Dup1_2F | TTTTGCGGTCCATGCACGAT |
| Coeaexf_Dup1_2R | GAGCCGTCGAAGATGTCCTT |
| Coeaexf_Dup1_2Rc1 | GCTTTTCCAGCGTTTCCAGC |
| Coeaexf_Dup2_2F | AATGTACCCGTTCAGCAGCT |
| Coeaexf_Dup2_2R | CGGCAGATGTTACCACCGAA |
| Coeaexf_Dup2_2Rc1 | TGTGCAGCACTATCTGGAGG |

---

For all primer sets, cycling conditions are:

| | | |
|---|---|---|
| 94°C | | 3mins |
| 35 cycles of: | | |
| | 94°C | 30s |
| | 60°C | 30s |
| | 72°C | 45s |
| 72°C | | 10mins |

---

**Coeaexf_Dup1_2** primers:
Expected CNV band size: 178
Expected control band size: 429
<u>per reaction:</u>

| | |
|---|---|
| water | 2.95ul |
| 10x PCR buffer | 1ul |
| 2mM dNTPs | 1ul |
| 5mM primer Coeaexf_Dup1_2F | 2ul |
| 5mM primer Coeaexf_Dup1_2R | 1ul |
| 5mM primer Coeaexf_Dup1_2Rc1 | 1ul |
| Taq | 0.05ul |
| DNA | 1ul |
| Total | 10ul |

**Coeaexf_Dup2_2** primers:
Expected CNV band size: 167
Expected control band size: 400
<u>per reaction:</u>

| | |
|---|---|
| water | 2.95ul |
| 10x PCR buffer | 1ul |
| 2mM dNTPs | 1ul |
| 5mM primer Coeaexf_Dup2_2F | 2ul |
| 5mM primer Coeaexf_Dup2_2R | 1ul |
| 5mM primer Coeaexf_Dup2_2Rc1 | 1ul |
| Taq | 0.05ul |
| DNA | 1ul |
| Total | 10ul |

#### **Supplementary Figure 5** - CNV primer PCR validation

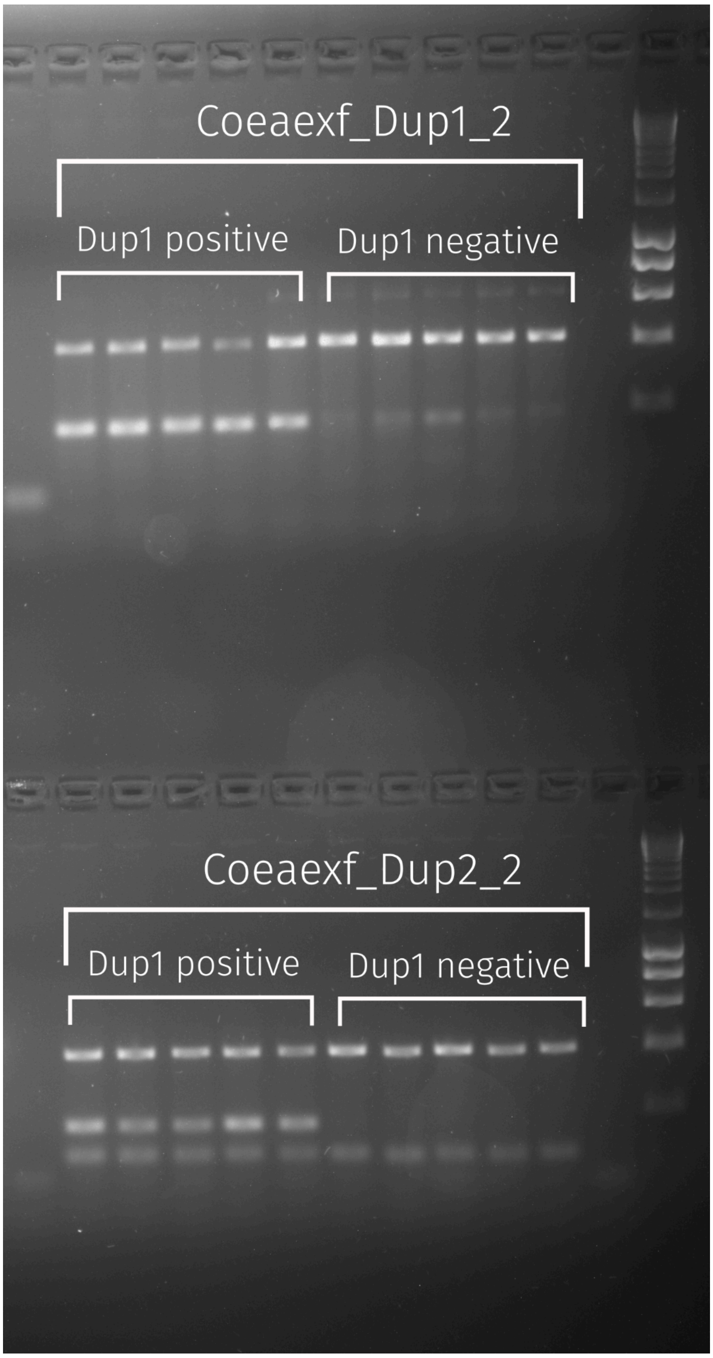


**Supplementary Figure 5.** A gel image of the two duplication diagnostic PCR primers, applied to *An. gambiae* (Dup1) and *An. arabiensis* (Dup2) individuals. Five Dup-negative samples and five Dup-positive samples were tested for each primer pair. Concordance is 100%.
